# Bacteria with antibacterial activities isolated from *Magallana gigas* microbiota as potential probiotics against *Vibrio aestuarianus* infections in oyster farming

**DOI:** 10.1101/2025.04.07.647571

**Authors:** Luc Dantan, Marie-Agnès Travers, Lionel Degremont, Benjamin Morga, Prunelle Carcassonne, Mickael Mege, Yannick Fleury, Bruno Petton, Elise Maurouard, Jean-François Allienne, Gaëlle Courtay, Océane Romatif, Raphaël Lami, Laurent Intertaglia, Yannick Gueguen, Jeremie Vidal-Dupiol, Céline Cosseau, Eve Toulza

**Affiliations:** IHPE, Univ Perpignan Via Domitia, CNRS, IFREMER, Univ Montpellier, Perpignan, France; IHPE, Univ Montpellier, CNRS, IFREMER, Univ Perpignan Via Domitia, Montpellier, France; Ifremer, ASIM, F-17390 La Tremblade, France; Laboratoire de Biotechnologie et Chimie Marine, EA3884, Université de Bretagne Occidentale, Université Bretagne Sud, 29334 Quimper, France; Ifremer, UBO CNRS IRD, LEMAR UMR 6539 Argenton, France; Sorbonne Université, UPVD, CNRS, Laboratoire de Biodiversité et Biotechnologies Microbiennes, F-66650 Banyuls-sur-Mer, France; Sorbonne Université, CNRS, Bio2Mar, Observatoire Océanologique, 66650 Banyuls-sur-Mer, France; MARBEC, Univ Montpellier, CNRS, Ifremer, IRD, Sète, France

**Keywords:** *Magallana gigas*, Antibacterial activities, Microbiota, *Vibrio aestuarianus*, Aquaculture

## Abstract

**Introduction:** Oyster farming is a significant industry worldwide, but it is threatened by various diseases such as Pacific Oyster Mortality Syndrome or vibriosis. *V. aestuarianus* is a major cause of mortality for market-size oysters, resulting in significant economic losses for oyster farmers. Among the various control methods developed, probiotics appear to be a promising approach. More specifically, the use of the antibacterial activity of bacteria from the natural microbiota of the oyster *Magallana gigas* appears to be a sustainable solution against *V. aestuarianus* infections.

**Results:** Our study investigated the probiotic potential of bacteria isolated from the microbiota of *M. gigas* oysters. We screened a collection of 334 bacteria against eight target pathogens, including *V. aestuarianus*, and identified 78 bacteria with antibacterial activity for which eight retained this activity in their culture supernatants. Five strains were selected for further testing and exposed to oysters prior to *V. aestuarianus* infection. Our results show that four strains significantly reduced oyster mortality, with a maximum reduction of 70%. In addition, changes in oyster microbiota composition were observed following exposure, but the administered bacteria were not detected in the microbiota.

**Conclusion:** Our findings demonstrate the potential of oyster microbiota-derived bacteria as probiotics for disease control in oyster farming. This approach could provide a sustainable and environmentally friendly solution for the oyster farming industry. Further research is needed to understand the underlying mechanisms and to develop effective probiotic-based strategies for preventing *V. aestuarianus* infection.

## Introduction

The Pacific Oyster *Magallana gigas* (formerly known as *Crassostrea gigas*), is the most widely cultivated oyster species in the world, contributing significantly to the aquaculture industry (Food and Agriculture Organisation 2022). Nevertheless, the farming of *M. gigas* encounters substantial difficulties due to recurrent infectious diseases, leading to high annual mortality rates (Friedman et al. 2005; Cotter et al. 2010; Pernet et al. 2012; Azéma et al. 2015). Since 2001, mass mortality of adult *M. gigas* has been reported in France, in association with the detection of the bacterium *Vibrio aestuarianus* (Garnier et al. 2008). This bacterium is a harmful primary pathogen with chronic mortality reaching a cumulative mortality rate up to 30%. This represents important economic consequences since *V. aestuarianus* preferentially infects market size oysters which have been raised for several years (Azéma et al. 2017; Lupo et al. 2019). Other *Vibrio* species have been associated with mortality episodes affecting *M. gigas* oysters at different stages of development. Notably, *V. coralliilyticus* has been linked to massive mortalities of *M. gigas* larvae (Richards et al. 2015; Travers et al. 2015; Ushijima et al. 2022). Spat / juveniles are affected by *V. crassostreae* (Dégremont et al. 2021; Cowan et al. 2023) and *V. harveyi* (Dégremont et al. 2021; Oyanedel et al. 2023).

Efforts to address the challenges of *Vibrio* infections affecting *M. gigas*, have led to various strategies build upon growing knowledge on oysters. One promising method involves the use of genetic selection to breed pathogen-resistant oysters (Dégremont et al., 2015, 2020), although this approach has limitations, such as the potential selection of trade-offs that could negatively impact the commercial value of *M. gigas*. Furthermore, the discovery of immune priming in *M. gigas* has paved the way for innovative applications using heat-killed *V. splendidus* (Zhang et al. 2014), which provide protection against *V. splendidus* infections. Research on disease prevention in molluscs based on the use of probiotics has been ongoing for decades (Yeh et al. 2020; Takyi et al. 2023, 2024; Dantan et al. 2024a, b; Muñoz-Cerro et al. 2024). Probiotics display their positive benefits through a variety of methods, including direct pathogen inhibition via competition for nutriments or production of antimicrobial compounds, but also indirect immunomodulatory effects (Lazado and Caipang 2014; Yan et al. 2014; Peixoto et al. 2017; Khademzade et al. 2020). Previous studies have demonstrated that bacterial strains *Pseudoalteromonas sp.* hCg-6 and *Pseudoalteromonas sp.* hCg-42 isolated from *M. gigas* haemolymph, displayed *in vitro* antibacterial activity against marine pathogens such as *V. splendidus*, *V. tapetis*, *V. harveyi* and *Aeromonas salmonicida* (Defer et al. 2013; Desriac et al. 2014; Offret et al. 2018). Furthermore, exposure to *Pseudoalteromonas* sp. hCg-6 has been shown to enhance the survival of *Haliotis tuberculata* abalone during infection with *V. harveyi* ORM4 (Offret et al. 2018). In addition, antimicrobial-producing bacteria have been employed as a strategy to improve the survival of oysters against bacterial infections. For instance, *Crassostrea virginica* larvae exposed to *Bacillus pumilus* RI06-95 exhibited significantly increased survival rates during challenge with *Vibrio coralliilyticus* (Sohn et al. 2016). Similarly, an exposure of *M. gigas* larvae to *Pseudoalteromonas sp.* was found to inhibit the growth of *Vibrio coralliilyticus*, thereby improving larval survival during subsequent infection (Madison et al. 2022).

In this article, we investigated the potential of bacteria isolated from the natural microbiota of *M. gigas* to protect oysters against *V. aestuarianus* infection. For this, we firstly screened a collection of bacteria previously isolated from *M. gigas* associated microbiota (Dantan et al. 2024b) for their antibacterial activity *in vitro* against four oyster pathogenic Vibrio. sp. (Travers et al. 2015) and against opportunistic bacteria associated with the POMS disease (de Lorgeril et al. 2018; Clerissi et al. 2022). Secondly, selected candidate bacteria were tested for their effect against *V. aestuarianus* infection, and we investigated the impact of the administered bacteria on the microbiota of the exposed oysters.

## Materials and methods

### Screening for antibacterial activities of bacteria isolated from *M. gigas* microbiota

Eight bacterial strains were selected as targets for the screening of antibacterial activities. Four of them were pathogenic *Vibrio* for oysters at different developmental stages: *Vibrio aestuarianus* 02/041 (Garnier et al. 2008), *Vibrio coralliilyticus* 06/210 (Dégremont et al. 2021), *Vibrio crassostreae* J2-9 (Lemire et al. 2015) and *Vibrio harveyi* Th15_O_A01 (Oyanedel et al. 2023). The four others are bacteria associated with POMS dysbiosis according to (de Lorgeril et al. 2018; Clerissi et al. 2022): *Amphitrea sp.* 14/114-3T2, *Marinobacterium sp.* 05/091-3T1, *Marinomonas sp.* 12/107-2T2, *Pseudoalteromonas sp.* 09/041-1T3 (**Supplementary_File_1 Table S1**). Target bacteria were provided by the French National Reference Laboratory (Ifremer, La Tremblade, France) or came from previous projects carried out in our laboratory (Oyanedel et al. 2023).

All strains were cultivated from glycerol stock in 10 mL Marine Broth (MB) at 20°C for 48 hours under constant agitation (100 rpm), then the OD_600_ was determined using BioPhotometer (Eppendorf). The cultures were diluted into fresh MB to a final concentration of 10^6^ CFU/mL prior to inoculation of Marine Agar plates by inundation. The 334 bacteria from our previously described bacterial collection isolated from *M. gigas* microbiota (**Supplemenary_File_1 Table S2**) (Dantan et al. 2024b) were then tested for their antibacterial activity against each of the target. Each bacteria from the collection were grown from glycerol stock in 2 mL MB at 20°C under constant agitation (100 rpm) for 48h before being distributed onto 4 different 96 well microplates. These microplates were then duplicate using a microplate pin replicator on new 96 well microplate containing fresh MB and incubated overnight at 20°C on MB media under constant agitation (100 rpm) and then deposited in arrays of 8×12 (2 µL) spots using a microplate pin replicator on marine agar plate previously inoculated with the target bacteria. A 2 µL spot of kanamycin (50 µg/mL) was used as positive control and a 2 µL spot of sterile Marine Broth as a negative control. Marine agar plates without target bacteria were used as growth and purity control of the bacteria from collection. Agar plates were then incubated at 20°C for two days and were then photographed using Gel Doc XR (Biorad, CA, USA) and the “Flamingo” filter to visualise a potential halo of inhibition characteristic of antibacterial activity. For supernatant assay, the same target bacteria were used. Prior to the test, the bacteria were cultured in Marine Broth media on 96 well plates during 72h at 20°C. After the incubation period, the 96 well plate was centrifugated during 10 minutes at 4000 rpm. The supernatants were then carefully transferred into new 96 well plates and heated at 100°C for 5 minutes to kill the possible remaining bacteria. Then, 2 µL spots were deposited on the marine agar plates previously flooded with the target bacteria as describe above.

### Oysters used as donors and recipient

Oysters were produced at the Ifremer hatchery in La Tremblade in February 2021. Briefly, 25 females and 25 males were used to produce100 bi-parental families (each male was mated to four females, and each female was mated to four males).

Each family was raised in separated tank during the larval stage, and then each family was settled in separate trays until two-months old. Then, 150 spat of each family were individually counted and mixed together to produce a batch. This batch of mixed families was transferred to the Ifremer nursery in Bouin in May 2021 until the experimental infection in November 2021. All oysters were kept in our controlled facilities using UV-treated seawater until their evaluations. Animals were fed *ad libitum* using a cultured phytoplankton diet (*<u>Isochrysis</u> <u>galbana</u>*, *<u>Tetraselmis suecica</u>* and *<u>Skeletonema costatum</u>*).

### Oyster exposure to bacteria selected for a potential beneficial effect

Recipient oysters were distributed between seven 40 L tanks filled with UV-treated seawater and maintained at 20°C with adequate aeration. Each tank contained 75 adult oysters (mean individual weight = 29.68 ± 8.03 g). These recipient oysters were either exposed to one of the five bacterial strains selected for their antibacterial activities, one control bacteriocin producing strain (*Pseudoalteromonas sp.* hCg-42) which has been previously isolated from *M. gigas* haemolymph (Defer et al. 2013) or to sterile artificial seawater (control).

Prior to exposure, the bacteria (*Pseudoalteromonas* sp. hCg-42, *Bacillus* sp. ARG61, *Halomonas* sp. LTB66, *Cytobacillus* sp. ARC29, *Yoonia* asp. THAU59 and *Vibrio* sp. LTB1) were individually cultured in 10 mL of MB media for 48h at 20°C under constant agitation and then, 1 mL of each bacterial culture was inoculated into 10 mL fresh MB media and incubated at 20°C under constant agitation. After 48 hours of incubation, the OD_600_ was measured, and the appropriate amount of bacteria (1 OD_600_ unit = 8×10^8^ CFU/mL) was collected, centrifuged at 4000 rpm for 2 minutes and the supernatant was discarded. The pellets were then resuspended in 10 mL sterile artificial seawater and added immediately to tanks containing the adult oysters so that the final concentration in the tank was adjusted to 10^4^ CFU/mL. The selected bacteria were added individually during seven days and were renewed two times without water renewal at days two and four (**Figure 1**).

**Figure 1:**
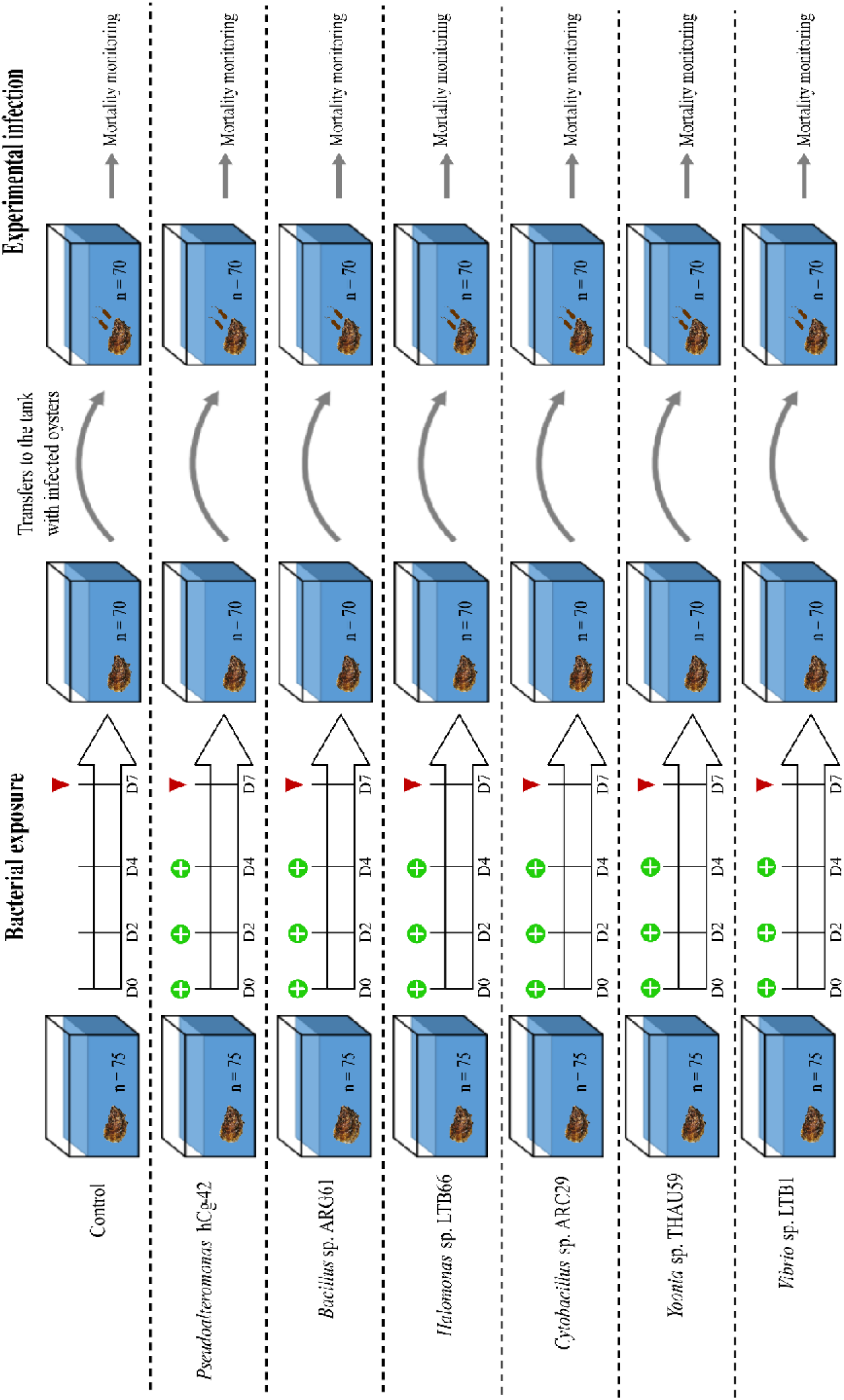
Overall experimental design for bacterial exposure and experimental infections performed with the NTA oyster population. Recipient oysters (n=75) were placed in 40L tank filed with UV-treated seawater and maintained at 20°C with adequate aeration. Oysters were then exposed to individual selected bacterial strains during seven days with a renewal every two days (indicated by the “+” sign in the green circle). At the end of bacterial exposure, 5 oysters per tank were sampled, flash-frozen into liquid nitrogen and stored at-80°C for molecular analysis (indicated by red triangles). Right after the bacterial exposure, remaining recipient oysters were transferred into 5 new tanks containing donor oysters injected with *V. aestuarianus* in order to realise an experimental infection.

Five oysters exposed to each condition were sampled at day seven (D7) of the bacterial exposure to perform molecular analysis. Sampled oysters were grounded in liquid nitrogen (Retsch MM400 mill) to a powder that was stored at-80°C until subsequent DNA extraction.

### *Vibrio aestuarianus* experimental infection by cohabitation

A *V. aestuarianus* experimental infection was performed immediately following the bacterial exposure to recipient oysters. A cohabitation protocol was used as previously described in (De Decker and Saulnier 2011). The *V. aestuarianus* 02/041 was grown in Zobell medium at 22°C for 24h under agitation. The bacterial concentration was determined by measuring the OD_600_ and was adjusted to OD_600_ of 1 representing 5.10^8^ bacteria per mL in artificial seawater. Seventy donor oysters were injected in the adductor muscle with 100µL of the *V. aestuarianus* 02/041 suspension and were then equally distributed among the five tanks (**Figure 1**). The donors were from the same oyster population as the recipient oysters exposed to the strains showing antibacterial activities. A ratio 1:1 was used for donor and recipient oysters (Azéma et al. 2017). After 48 hours of cohabitation and before first mortality, donors were removed from the tanks. The mortality of recipient oysters exposed to selected bacteria and control oysters, was then recorded during 17 days by recording the dead oyster every day, and all the dead oysters were removed from the tanks.

### Statistical Analysis of oyster mortality

Oyster mortality was analysed using survival analysis performed on R (v 4.2.1) (R Core Team 2022) with the package survminer (v 0.4.9) (https://cran.r-project.org/web/packages/survminer/index.html). The Kaplan-Meier method was used to represent the cumulative survival rate. A log-rank test was used to determine the difference between the conditions and post-hoc pairwise comparisons with Bonferroni corrected p-value were used to define which values were significantly different from the control. A multivariate Cox proportional hazards regression model was used to compute Hazard-Ratio (HR) with confidence intervals of 95%.

### Bacteria and oyster DNA extraction

DNA extraction from oysters collected during bacterial exposure was performed from frozen powders using DNA from the tissue Macherey-Nagel kit according to the manufacturer’s protocol. Prior to 90 min of enzymatic lysis in the presence of proteinase K, an additional 12-min mechanical lysis (Retsch MM400 mill) was performed with zirconia/silica beads (BioSpec). DNA concentrations were checked with a Qubit® 2.0 Fluorometer (Thermo Scientific) and adjusted when necessary.

Bacterial DNA for the candidate strains used to constitute the mock community was extracted as described in (Dantan et al. 2024b)

### 16S rDNA library construction and sequencing

Library construction (with primers 341F 5’-CCTAYGGGRBGCASCAG and 806R 5’-GGACTACNNGGGTATCTAAT targeting the 16S V3V4 region) and sequencing on a MiSeq v2 (2×250 bp) were performed by ADNid (France).

### Bioinformatic pipeline for 16S barcoding analysis

Sequencing data obtained in this study were processed with the SAMBA (v 3.0.2) workflow developed by the SeBiMER (Ifremer’s Bioinformatics Core Facility). Briefly, Amplicon Sequence Variants (ASV) are constructed with DADA2 (Callahan et al. 2016) and the QIIME2 dbOTU3 (v 2020.2) tools (Bolyen et al. 2019), Due to the known diversity overestimation generated by DADA2, an additional step of ASV clustering has been performed using dbOTU3 algorithm (Olesen et al. 2017) and contaminations were removed with microDecon (v 1.0.2) (McKnight et al. 2019). Taxonomic assignment of ASVs was performed using a Bayesian classifier trained with the Silva database v.138 using the QIIME feature classifier (Wang et al. 2007). Finally, community analysis and statistics were performed on R (R version 4.2.1) (R Core Team 2022) using the packages phyloseq (v 1.40.0) (McMurdie and Holmes 2013), Vegan (v 2.6-4) (Oksanen et al. 2022) and MicroEco (v. 1.9.1) (Liu et al. 2021).

For beta-diversity, the ASVs counts were preliminary normalized with the “rarefy_even_depth” function (rngseed = 711) from the package phyloseq (v 1.40.0)(McMurdie and Holmes 2013). Principal Coordinates Analysis (PCoA) were computed to represent dissimilarities between the samples using the Bray-Curtis distance matrix. Differences between groups were assessed by statistical analyses (Permutational Multivariate Analysis of Variance) using the adonis2 function implemented in the vegan package (2.6-4) (Oksanen et al. 2022).

### Detection of administered bacteria in 16S barcoding dataset

In order to search for the specific presence of the administered bacteria, we first produced full-length 16S DNA for the identification of the selected candidate bacteria (Dantan et al. 2024b). In parallel, a mock community composed of equal amounts of DNA from four of the administered bacteria (*Bacillus* sp. ARG61, *Vibrio* sp. LTB1, *Halomonas* sp. LTB66, and *Cytobacillus* sp. ARC29) was also submitted to 16S amplicon sequencing in order to validate our method. We then aligned 16S reference sequences of the administered bacteria against all the ASV sequences from the dataset using BLAST (Altschul et al. 1990). We considered ASVs sequences with a percentage of identity superior to 99% along the full V3V4 marker sequence as being our administered bacteria.

## Results

### 78 bacterial strains displayed an antibacterial activity against

We previously isolated 334 bacteria from the microbiota of *M. gigas* oysters from four different geographical sites (Brest Bay, La Tremblade in Marennes-Oleron Bay, Arcachon Bay and Thau Lagoon) (Dantan et al. 2024b). This collection was screened for their antibacterial activity against eight target bacteria (**Supplementarry_File_1_Table S1**). Among the 334 bacteria from the collection, 78 strains showed an inhibition area around bacterial colony. Among these strains, 32 showed antibacterial activity against Vibrios (17 against *V. harveyi* Th15_O_A01, 14 against *V. aestuarianus* 02/041, 12 against *V. coralliilyticus* 06/210 and 8 against *V. crassostreae* J2-9) and 65 against opportunistic bacteria associated with POMS disease (49 against *Marinomonas sp.* 12/107-2T2, 8 against *Pseudoalteromonas sp.* 09/041-1T3, 6 against *Amphitrea sp.* 14/114-3T2, and 2 against *Marinobacterium sp.* 05-091-3T1) (**Figure 2A**).

**Figure 2:**
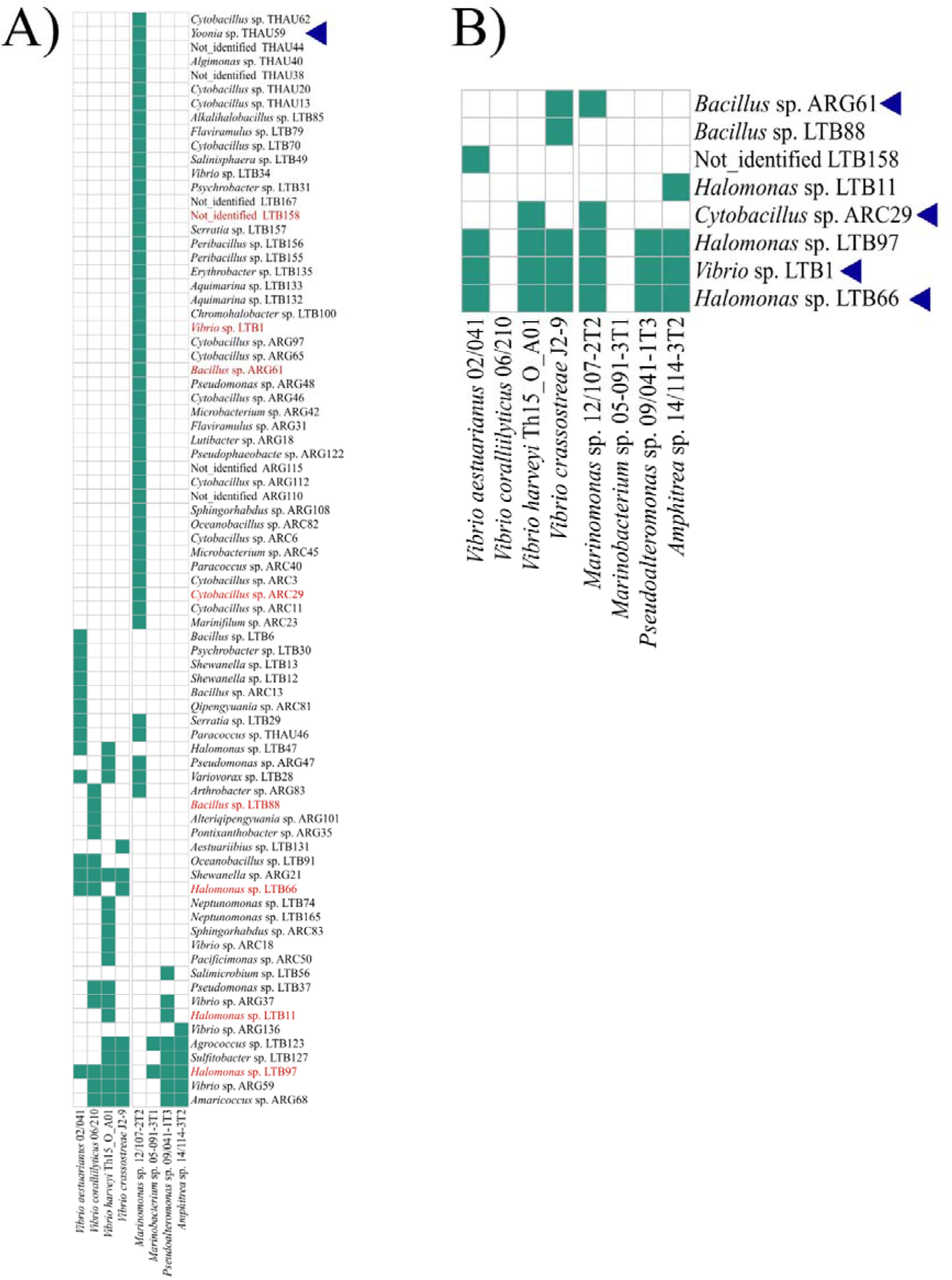
Antibacterial activities of bacteria isolated from oysters against the eight target bacteria. The plot represents all the bacteria of the collection having an antibacterial activity against the different target bacteria (A) by coculture and (B) in the culture supernatant. Positive antibacterial activity tests are represented by green tiles. Bacteria written in red (A) are those for which activity was found in the culture supernatants (B). Bacteria selected for experimental infection assay are indicated by a blue triangle.

The 76 bacteria presenting an antibacterial activity by co-culture were further screened for the antibacterial activity of their culture supernatant. Thus, height bacteria conserved their activity in the culture supernatant. Among them, three bacterial strains (*Halomonas sp.* LTB66, *Halomonas sp.* LTB97 and *Vibrio sp.* LTB1) presented antibacterial activities in their supernatant against six of the eight target bacteria, two strains (*Bacillus sp.* ARG61 and *Cytobacillus sp.* ARC29) presented antibacterial activities in their culture supernatant against two of the height target bacteria and the three other strains had antibacterial activities in their supernatant against one of the eight target bacteria. (**Figure 2B**).

Based on these results, we selected five bacterial strains for further assay. The bacteria were chosen to ensure representation from each geographic origin in the selected bacteria, preferably with antibacterial activity present in the supernatant culture. Thus, *Bacillus sp.* ARG61 from Brest Bay, *Halomonas sp.* LTB66 and *Vibrio sp.* LTB1 from La Tremblade in Marennes-Oleron Bay, *Cytobacillus sp.* ARC29 from Arcachon Bay and *Yoonia sp.* THAU59 from Thau Lagoon were selected for protection assay during an experimental infection.

### Four bacterial strains induced a significant reduction of mortality risk during *V. aestuarianus* infection

To test if exposure to selected bacteria with an antibacterial activity can induce a protective effect against *V. aestuarianus* infection, adult oysters (exposed or control) were challenged with *V. aestuarianus*.

The first mortalities were observed 96 hours post cohabitation with donor oysters (**Figure 3**). Compared to control condition (survival_t=408h_ = 0.14), a significant increase in survival was observed at t = 408 hours post-cohabitation for oyster exposed to bacterial strains *Pseudoalteromonas sp.* hCg-42 (survival_=408h_ = 0.60; p-value < 0.001), *Yoonia sp.* THAU59 (survival_=408h_ = 0.49; p-value < 0.001), *Bacillus sp.* ARG61 (survival_=408h_ = 0.47; p-value = 0.004), and *Cytobacillus sp.* ARC29 (survival_=408h_ = 0.40; p-value = 0.031). No difference in survival was observed for oyster exposed to *Halomonas sp.* LTB66 (survival_=408h_ = 0.19; p-value = 1) or *Vibrio sp.* LTB1 (survival_=408h_ = 0.07; p-value = 1) (**Figure 3**)(**Table 1**). A forest plot analysis confirmed these results and indicates that a significant reduction of the mortality risk of 70% (Log-Rank test: p-value < 0.001), 54% (Log-Rank test: p-value < 0.001), 46% (Log-Rank test: p-value = 0.002) and 58% (Log-Rank test: p-value < 0.01) was observed for the adult oysters exposed to the bacterial strains *Pseudoalteromoans sp.* hCg-42, *Bacillus sp.* ARG61, *Cytobacillus sp.* ARC29 and *Yoonia sp.* THAU59 respectively (**Supplementary_File_2 Figure S1**).

**Figure 3:**
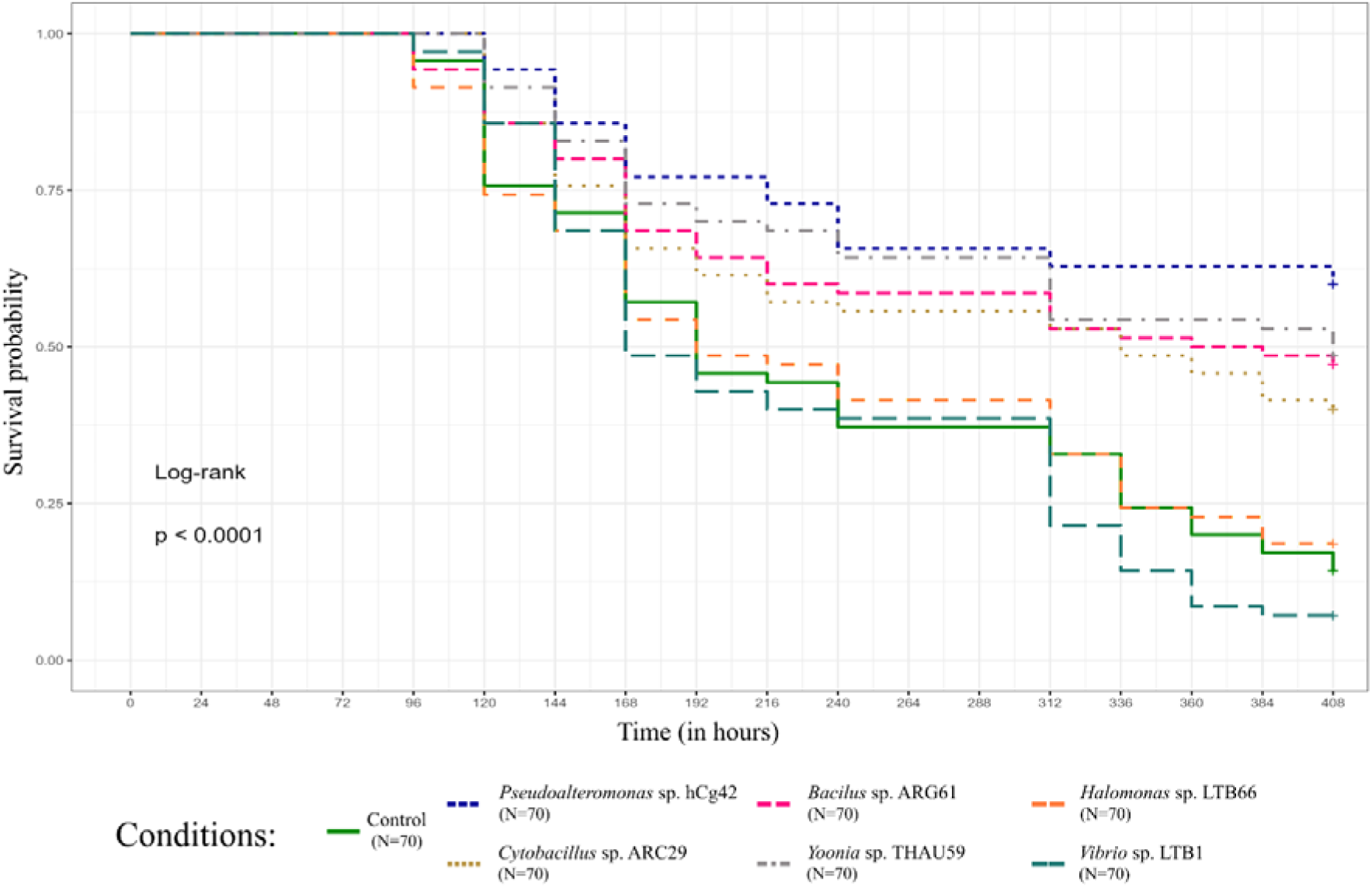
Four bacterial strains have improved oyster survival against *V. aestuarianus* infection. Kaplan-Meier curve representing survival probability of oysters for the control (green solid line) or exposed to candidate bacteria conditions during *Vibrio aestuarianus* infection.

**Table 1:** Oyster survival data at t=408h post *V. aestuarianus* infection.

| Condition | survival<br>(at t=408h) | std.err | p-value |
| --- | --- | --- | --- |
| Control | 0.143 | 0.042 | NA |
| *Pseudoalteromonas* sp. hCg-42 | 0.600 | 0.059 | 0.000001 |
| *Bacillus* sp. ARG61 | 0.471 | 0.060 | 0.00398 |
| *Halomonas* sp. LTB66 | 0.186 | 0.047 | 1.00000 |
| *Cytobacillus* sp. ARC29 | 0.400 | 0.059 | 0.03107 |
| *Yoonia* sp. THAU59 | 0.486 | 0.060 | 0.00032 |
| *Vibrio* sp. LTB1 | 0.071 | 0.031 | 1.00000 |

### Addition of beneficial strains slightly modifies the bacterial alpha diversity of the oyster microbiota

To test the immediate effect of the bacterial exposure on oyster microbiota, we analysed the bacterial communities by 16S rRNA gene sequencing after seven days of bacterial exposure with the last addition of bacteria performed 72 hours before sampling (**Figure 1**).

Sequencing of the V3-V4 hypervariable region of the 16S rRNA gene resulted in a total of 1,713,283 clusters for a total of 40 samples. After a quality control (deleting primers and low-quality sequences, merging, and removing chimeras) and ASV clustering, 1,268,467 reads (74%) with an average of 31,712 reads per sample were retained for downstream analyses.

Analysis of the alpha diversity metrics (richness, Shannon and Pielou) (**Figure 4**) indicates that there are no significant differences for the species richness between the different conditions of bacterial exposure (**Figure 4A**). However, we observed a greater range of diversity in the oyster microbiota from the control (217 – 502), and those exposed to *Pseudoalteromonas sp.* hCg-42 (82 – 570) and to *Bacillus sp.* ARG61 (94 – 590) compared to *Halomonas sp.* LTB66 (289 – 429), *Cytobacillus sp.* ARC29 (262 – 595), *Yoonia sp.* THAU59 (267 – 535) or *Vibrio sp.* LTB1 (183 – 414) exposure. The alpha diversity with Shannon index (**Figure 4B**), which considers taxon diversity and abundance, coupled with the Pielou index (**Figure 4C**) which considers the distribution of individuals within species, show trends (NS Kruskal-Wallis test) where oysters exposed to *Cytobacillus sp*. ARC29 (Shannon: 4.35 – 5.08; Pielou: 0.77 – 0.81) and *Yoonia sp.* THAU59 (Shannon: 4.29 – 5.47; Pielou: 0.77 – 0.87) have more diverse bacterial communities with a more equitable distribution of species compared to the control condition (Shannon: 3.65 – 4.88; Pielou: 0.68 – 0.80) or the condition exposed to *Pseudoalteromonas sp.* hCg-42 (Shannon: 2.86 – 5.22; Pielou: 0,64 – 0.82).

**Figure 3:**
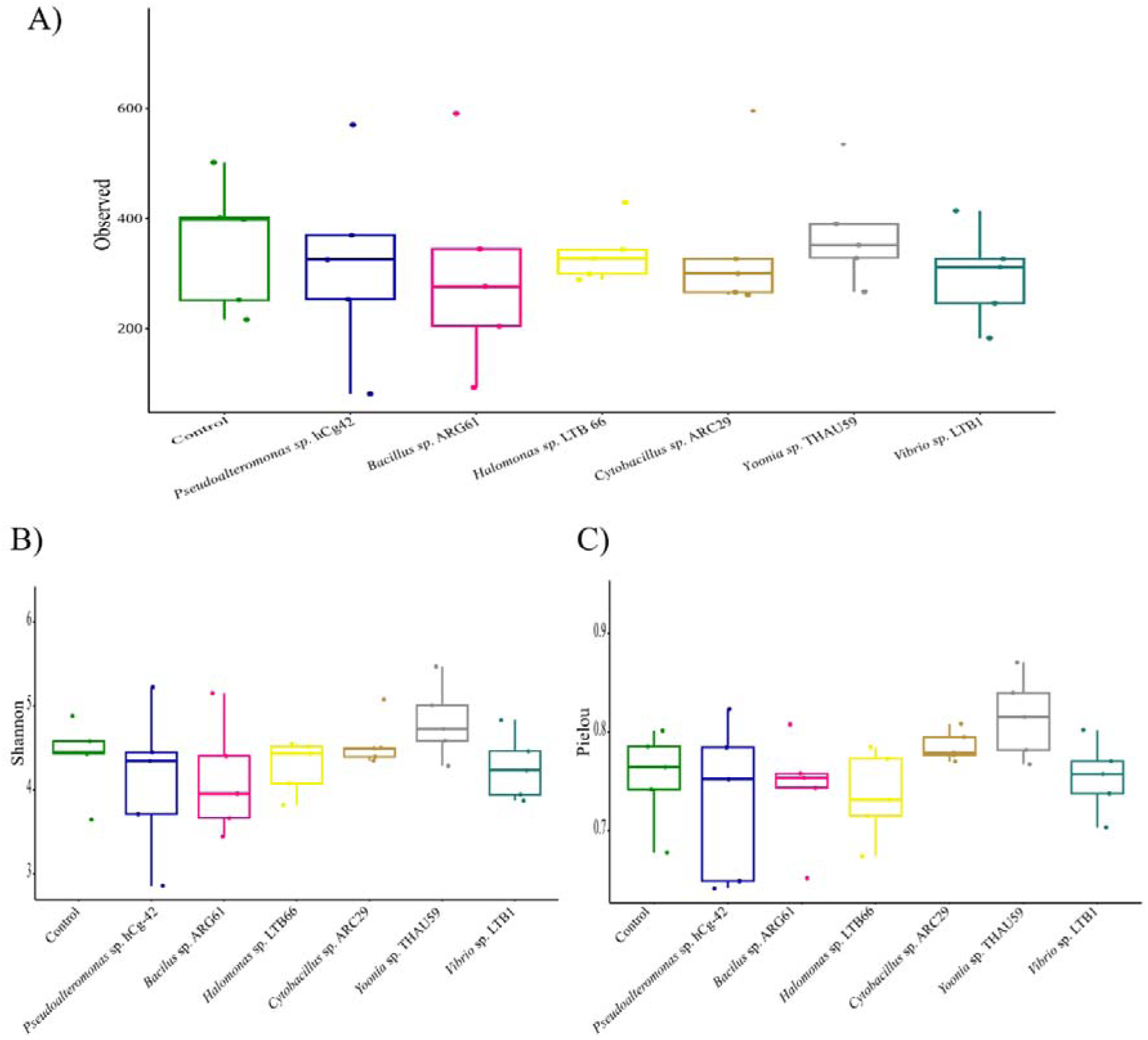
Seven days of bacterial exposure did not induce significant changes in alpha diversity. Boxplot representing the alpha diversity metrics (y axis) of oyster microbiota for control and exposed to selected bacteria (n=5 oyster per conditions) with Observed (A), Shannon (B) and Pielou (C) indices.

**Figure 4:**
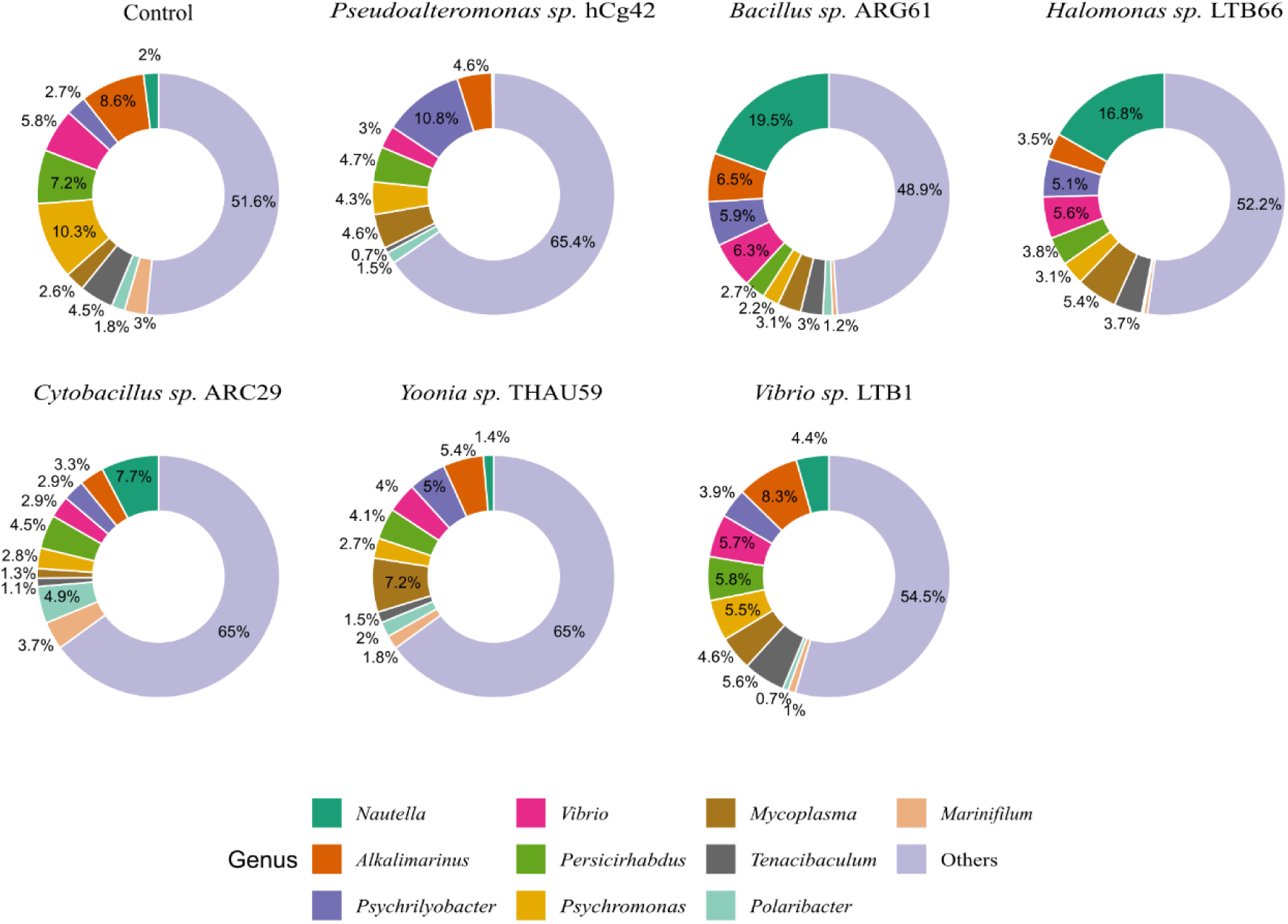
Bacterial composition differs according to bacterial exposure. Donut plot representing the mean relative abundance of bacterial communities for the control (n=5) or exposed to selected bacteria (n=5/condition) oyster samples at the genus level.

### Beneficial bacteria do not maintain in oyster tissues but significantly impact the beta diversity of the bacterial microbiota

After 7 days of exposure to the beneficial bacteria, we checked for their presence with the last addition of bacteria performed 72 hours before sampling. Doing a blast search against all the ASVs obtained using the 16S DNA sequences of the selected beneficial bacteria as a query, we were not able to detect the administered bacteria in the microbiota of oysters (**Supplementary_File_2 Figure S2A**) whereas the four bacteria were detected in the mock community although *Cytobacillus* and *Bacillus* were both affiliated to *Bacillus* (**Supplementary_File_2_Figure_S2A**). The ASVs from the Mock were thus manually reassigned to their bacterial strain in accordance with the BLAST results in particular for *Bacillus sp.* ARG61 and *Cytobacillus sp.* ARC29 (**Supplementary_File_2_Figure_S2B**).

Dissimilarity analysis, based on the Bray-Curtis index, on adult oyster microbiota revealed significant differences in microbiota composition between control oysters and oysters exposed to *Pseudoalteromonas* sp. hCg-42, *Bacillus* sp. ARG61, *Halomonas* sp. LTB66, *Cytobacillus* sp. ARC29, *Yoonia* sp. THAU59 and *Vibrio* sp. LTB1 (**Table 2**).

**Table 2:**
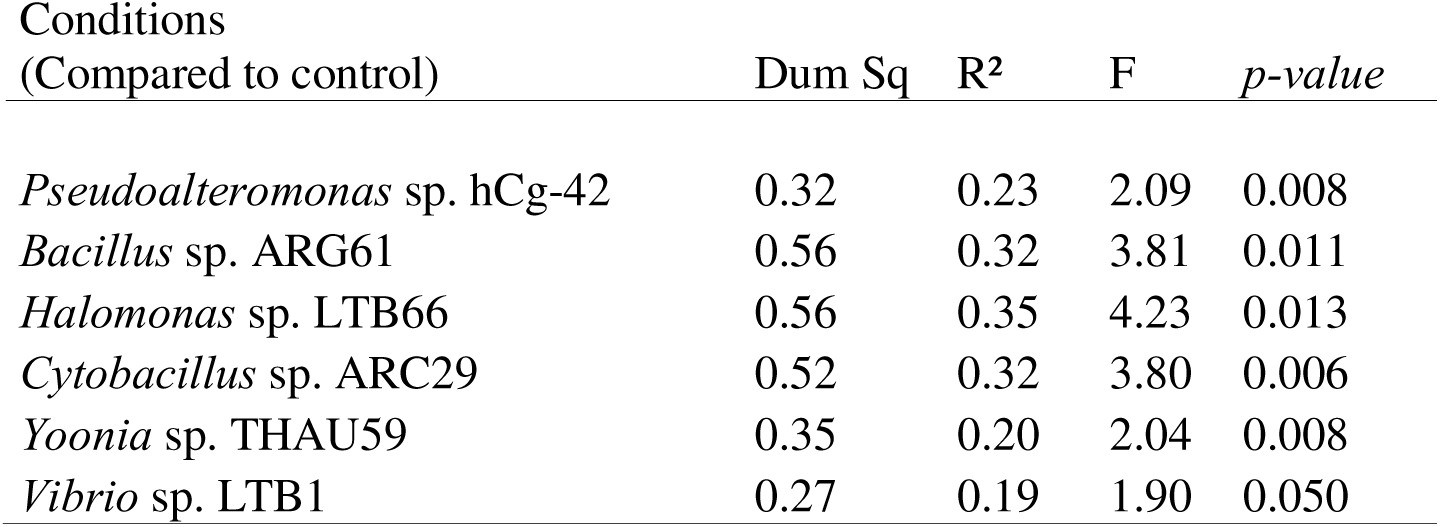
The beneficial bacterial exposure induces significant changes in the bacterial composition of the exposed oysters. Results of permanova on the Bray-Curtiss dissimilarity matrix showing the effects of microbial exposure on microbiota community composition compared to the control condition. Analyses were carried out on five oysters per condition, excepted for oysters exposed to Pseudoalteromonas sp. hCg-42, for which four oysters were used due to an abnormal sample that was discarded from the analysis. The p-values were obtained using 100,000 permutations.

At the phylum level, control oysters were dominated by Proteobacteria (54.1%) and Bacteroidota (23.2%) followed by Verrucomicrobiota (9.2%) and Firmicutes (3.6%) (**Figure 5A**). For oysters exposed to selected bacteria, Proteobacteria and Bacteroidota remained the dominant phyla. In comparison with the control condition, oysters exposed to *Pseudoalteromonas sp.* hCg-42 have a greater proportion of Firmicutes (13%) and Fusobacteriota (12.9%). The other conditions displayed a relatively similar composition compared to control condition (**Supplementary_File_2 Figure S3**). At the Genus level, in the control condition, the most abundant taxa were *Psychromonas* (10.3%), *Alkalimarinus* (8.6%), *Persicirhabdus* (7.2%) and *Vibrio* (5.8%) (**Figure 5**). In comparison, for oysters exposed to *Bacillus sp.* ARG61, *Halomonas sp.* LTB66 and *Cytobacillus sp.* ARC28 the most abundant genera were *Nautella* (19.5, 16.8 and 7.7% respectively), *Alkalimarinus* (6.5, 3.5 and 3.3%), *Psychrilyobacter* (5.9, 5.1 and 2.9%) and Vibrio (6.3, 5.6 and 2.9%). Oysters exposed to *Pseudoalteromonas sp.* hCg-42 displayed a higher proportion of *Psychrilyobacter* (10.8%) and *Mycoplasma* (4.6%). Oysters exposed to *Yoonia sp.* THAU59 displayed *Mycoplasma* (7.2%), *Alkalimarinus* (5.4%) and *Psychrilyobacter* (5%) as most abundant genera. Last for oysters exposed to *Vibrio sp.* LTB1 we identified *Alkalimarinus* (8.3%), *Persicirhabdus* (5.8%), *Vibrio* (5.7%) *Tenacibaculum* (5.6%) and *Psychromonas* (5.5%) (**Figure 5**).

## Discussion

Infectious diseases are a threat for oyster farming, and among them, the pathogenic bacterium *V. aestuarianus* has been observed to spread across Europe (Mesnil et al. 2022). To ensure the sustainability of oyster farming, it is crucial to develop effective, sustainable and socially accepted strategies to fight vibriosis. In this study, we explored the antimicrobial potential of bacteria isolated from oysters, with the aim of identifying natural antagonists that could mitigate the harmful effects of *V. aestuarianus* infections in oyster farms.

### Oyster microbiota is a promising source of bacteria with antibacterial activity

By screening a collection of 334 bacteria, we identified 76 bacteria (22.7% of the collection) that displayed an antibacterial activity by co-culture, and eight of them (2.4% of the collection) displaying a antibacterial activity in their supernatant against at least one of the target bacteria. We acknowledge that we may have lost heat sensitive antimicrobial molecules since our protocol to collect the supernatant include a heating step at 100°C. This could explain the small proportion of effective molecules present in the supernatant fractions. Furthermore, the spectrum of activity of the selected bacteria differed depending on if the antimicrobial tests were performed using coculture assay or using the culture supernatant. This could be explained by the release of an antibacterial compound during the centrifugation and/or the heating steps. It also underlines that the methodology used for the assay is a key issue for antimicrobial screening. Further tests using a different approach of supernatant extraction will therefore be necessary to be as exhaustive as possible. Furthermore, it will be interesting to carry out additional antibacterial activity tests using a broader range of pathogenic targets affecting species of aquaculture interest. This will help to decipher if the observed antimicrobial activity is restricted to oyster pathogens or if it can be expanded to applications to a broader range of species of economic interest.

### Exposure to four bacterial strains (*Pseudoalteromonas sp.* hCg-42; *Bacillus sp.* ARG61; *Cytobacillus sp.* ARC29 and *Yoonia sp.* THAU59) induced a significant reduction of the mortality risk against *V. aestuarianus* infection

Four bacterial strains (*Pseudoalteromonas sp.* hCg-42; *Bacillus sp.* ARG61; *Cytobacillus sp*.

ARC29 and *Yoonia sp.* THAU59) among 6 tested induced a significant reduction, from 46% to 70 %, of the mortality risk against *V. aestuarianus* infection. To our knowledge, this is the first demonstration of a protective effect against *V. aestuarianus* by a potential antagonistic bacterium. The Vibrio challenge was performed 72 hours after the last addition of the beneficial bacteria, and it will be important to test longer term effect. Indeed, the dynamics of Vibrio infection can be long and/or chronic-like (Travers et al. 2017). It is therefore possible that the protection we observed only delayed the progression of the infection.

Further analyses are also required to determine the mechanisms underlying resistance to *V. aestuarianus*. Here we did not detect the administered bacteria in the microbiota 72 hours after it was added. The absence of detection suggests that the added bacteria are likely not present as a major dominant strain in the microbiota. Our methodology was validated using a mock assay; however, we cannot rule out the possibility that the added strain may be present as a minority strain, which our method may have failed to detect due to a lack of sensitivity. Furthermore, this absence of detection is not surprising since it was reported several times in the literature that added probiotic strains do not maintain in their host microbiota. This can be observed in the European abalone *Haliotis tuberculata* where exposure to the *Pseudoalteromonas* hCg-6 exogenous strain, result in only a temporary presence of the probiotic strain probiotic strain in the haemolymph rather than the establishment of a long-term interaction (Offret et al. 2018). Exposure of *M. gigas* larvae to bacteriocin-like inhibitory substance (BLIS)-producing *Aeromonas*, showed that the probiotic strain concentration decreased right after it was added to the oyster and was not detectable 72 h after its addition (Gibson et al. 1998). Furthermore, among the four strains conferring a protective effect against *V. aestuarianus*, three bacteria *(Bacillus sp.* ARG61; *Cytobacillus sp.* ARC29 and *Yoonia sp.* THAU59) had no antibacterial activity against *V. aestuarianus in vitro*, making the explanation of resistance acquisition even more complex. Taken together, these results suggest that a direct antagonistic effect of the beneficial strains against *Vibrio* may likely not be at the origin of the resistance that we observe in our experiments. The change in the microbiota composition could rather explain the observed beneficial effect. Such an impact on microbiota composition after exposure to beneficial strains has been reported in other species. This beneficial shift in microbiota composition following an exposure to microorganisms has already been observed in Pacific oysters, where exposure to a complete microbiota or mixes of cultivable bacteria led to changes in microbiota coupled with improved survival against POMS infection (Fallet et al. 2022; Dantan et al. 2024b). This was also observed for *Crassostrea virginica* oyster larvae, which, following an exposure to the beneficial marine bacterium *Phaeobacter inhibens* S4, saw the composition of their microbiota modified, favouring in particular bacteria of the *Alteromonas* and *Pseudomonas* genus (Takyi et al. 2024). Exposure of *C. sikamea* oysters to *Streptomyces* strains N7 and RL8 also induced a shift in the diversity and composition of their microbiota, leading to a decrease in *Vibrio* bacteria (García Bernal et al. 2017). An alternative still non-exclusive hypothesis is that the exposure to these bacteria has induced immunostimulation effects as it has been previously reported in oysters, abalone and scallops. Specifically, studies have demonstrated that an exposure of *C. virginica* larvae to probiotic bacteria *Bacillus pumilus* RI06–95 and *Phaeobacter inhibens* S4 led to improved survival in the face of *V. coralliilyticus* RE22 infection simultaneously with an activation of immune signalling pathways (NF-kB and MAPK pathways) and expression of immune effectors such as serine protease inhibitor (Cv-spi2), mucins and antimicrobial histone H2B (Modak and Gomez-Chiarri 2020). Similarly, an exposure of *M. gigas* larvae to a mix of probiotics bacteria, led to an increased expression of immune signalling proteins (TOLL) and immune effectors such as interleukin IL-17 or MD88. Furthermore, larvae exposed to this mix of probiotics showed an increased survival against *V. coralliilyticus* infection (Hesser et al. 2024). Comparatively, *Argopecten purpuratus* scallop larvae exposed to bacterial strains belonging to *Psychrobacter*, *Hydrogenophaga*, and *Shewanella* genera displayed no mortality after 24h post infection to *V. bivalvicida* VPAP30. This beneficial effect could be due to an immunomodulation of genes coding for opsonin, superoxide dismutase (SOD), Toll-like receptor (TLR) or Lysozyme following the bacterial exposure (Muñoz-Cerro et al. 2024). At last, New Zealand black-footed abalone *Haliotis iris*, displayed a significantly increase in the number of total haemocytes count (HTC), non-apoptotic cells, an higher percentage of ROS-positive cells and an higher viability following an exposure to multi-strain probiotics (Grandiosa et al. 2018).

## Conclusion

In this study, we have shown that the administration of bacteria isolated from *M. gigas* microbiota and displaying *in vitro* antibacterial activities against various pathogens or opportunists, could increase the survival of oysters against *V. aestuarianus* infectious challenges. Further studies will be needed to understand the molecular mechanisms involved in the tolerance conferred by these bacteria. It would also be interesting to characterise and purify molecules that are active against pathogens and opportunists. These molecules may offer a promising way to mitigate the effects of these infectious diseases. A sustainable oyster farming is a key to develop this growing industry in a context of global changes and emerging infectious diseases. The development of prophylactic methods such as the use of probiotics is a necessity.

## Acknowledgements

The authors thank Céline Garcia from Ifremer, EU Reference Laboratory for mollusc diseases (La Tremblade, France) for providing oyster bacterial isolates used as target bacteria. We are grateful to Leo Duperret, Emily Kunselman, Nicole Faury, Cyrielle, Lecadet and Delphine Tourbiez for their help during the oyster experimental infections. We also like to thank Antoine Jourdan and both hatchery and nursery teams at PMMLT La Tremblade and PMM Bouin for the supply and care of the oysters. We are grateful to the BIO2MAR platform (http://bio2mar.obs-banyuls.fr) for access to the instrumentation.

## Authors’ contributions

LDa, LDé, BM, BP, EM, GC and JVD contributed to oyster sampling. LDa, PC, YF, RL and LI contributed to bacteria collection. LDa, and PC performed antibacterial activities tests. LDa, LDé, BM, BP, and MM performed oyster experiments. LDa, JFA, CG, OR, JVD, CC and ET prepared samples and performed DNA extraction from the oyster samples for analyses. LDa, CC and ET performed microbiota analyses. LDa, LDé, BM, MAT, BP, MM, YF, YG, JVD, CC and ET conceptualized and designed the experiments. LDa, Xx, Xx, CC and ET wrote the original draft. LDa, YG, JVD, CC and ET were involved in funds acquisition. All authors have read and approved the final manuscript.

## Availability of data and materials

Raw sequence data for 16S sequencing for metabarcoding analysis have been made available through the SRA database (BioProject accession number PRJNA1183305, Link for reviewer: ***)

## Ethical approval

The animal (oyster *Magallana gigas*) testing followed all European regulations concerning animal experimentation. The authors declare that the use of genetic resources fulfils the French and EU regulations on the Nagoya Protocol on Access and Benefit-Sharing (French legislation 2019-486).

