## Supplementary_File_2 Figure S1-S3 for "Bacteria with antibacterial activities isolated from *Magallana gigas* microbiota as potential probiotics against *Vibrio aestuarianus* infections in oyster farming"

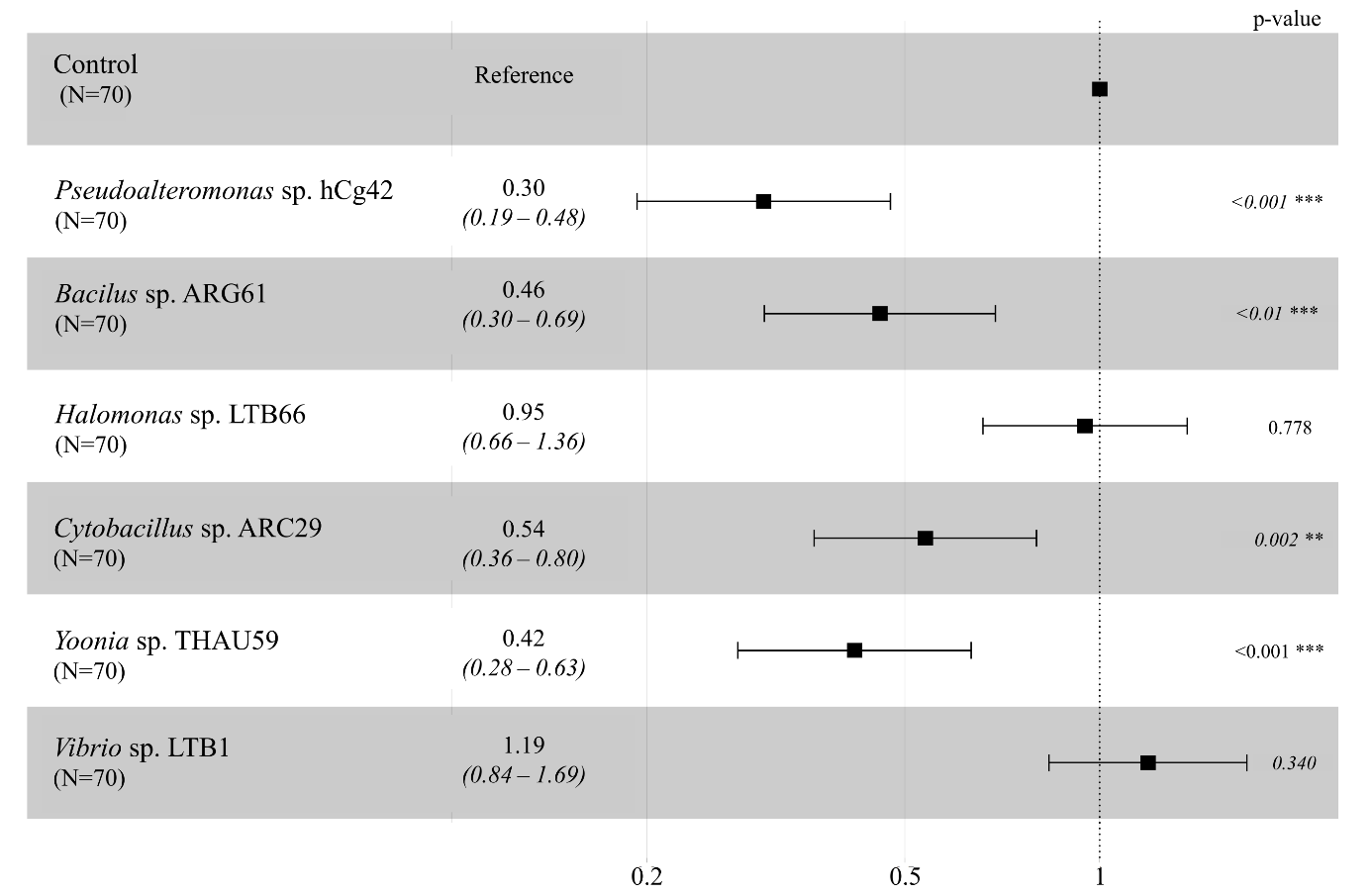

**Figure S1: Exposure to four bacterial strains has reduce the risk of mortality during a *V. aestuarianus* experimental infection**

Forest plot representing the Hazard-Ratio value of mortality risk during the *V. aestuarianus* experimental infection for oysters exposed to candidate bacteria compared to control oysters. The numbers in brackets under the different conditions correspond to the number of oysters used for the experimental infection. The Hazard-Ratio value is indicated to the right of the conditions, excepted for the control condition, which is indicated as reference. Finally, the value of the p-value is indicated on the right-hand side of each row.

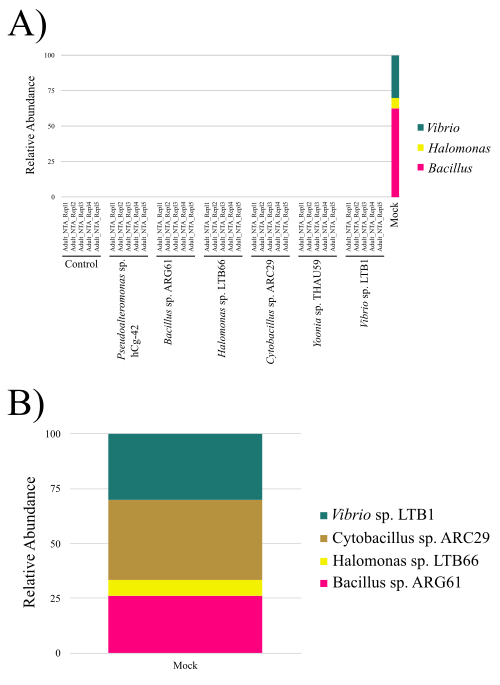

**Figure S2: The added bacteria were not found in the microbiota of oysters after seven days of exposure**

(A) Relative abundances, at the genus level, of the administered bacteria found after seven days of exposure in the oyster samples and for bacteria composing the mock control (*Bacillus* sp. ARG61, *Vibrio* sp. LTB1, *Halomonas* sp. LTB66 and *Cytobacillus* sp. ARC29). (B) Relative abundances, at the genus level, of bacteria composing the mock control (*Bacillus* sp. ARG61, *Vibrio* sp. LTB1, *Halomonas* sp. LTB66 and *Cytobacillus* sp. ARC29) after manually reassigned the genus according to BLAST result.

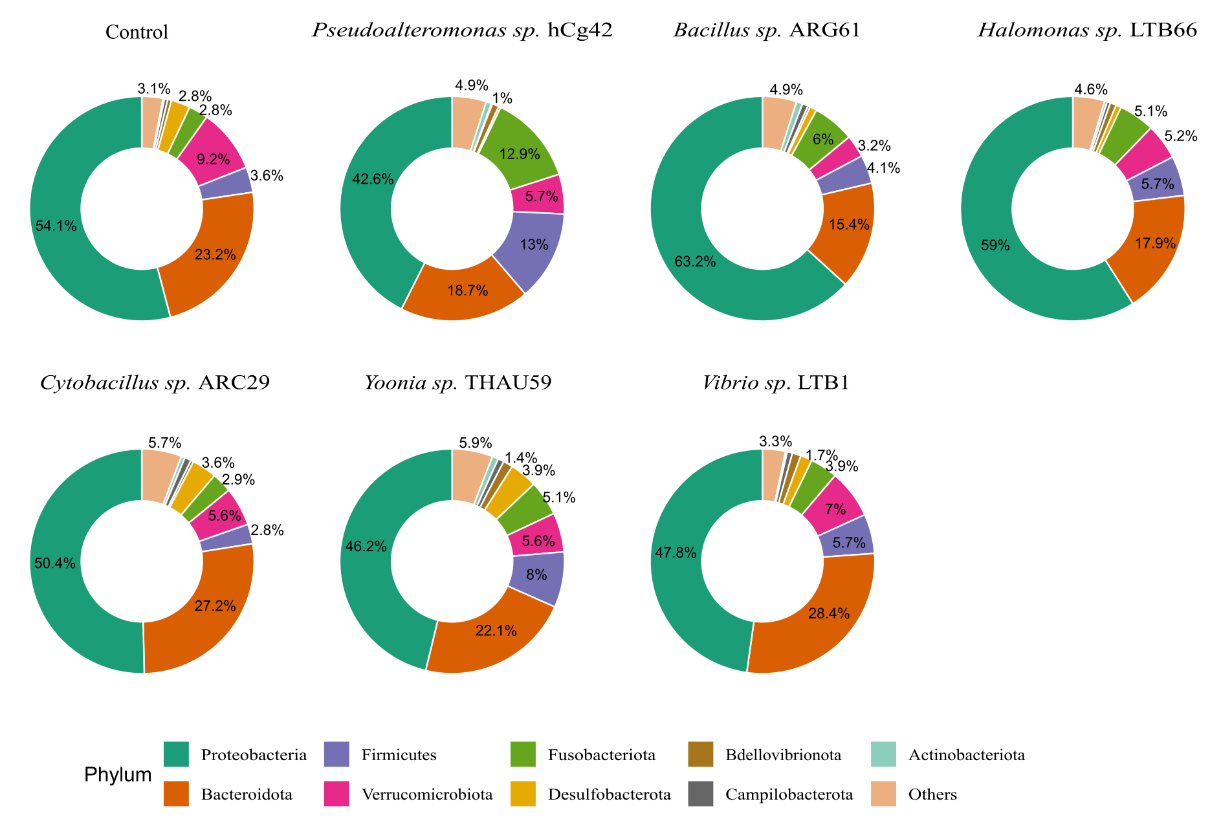

**Figure S3: Bacterial composition differs according to bacterial exposure.**

Donut plot representing the mean relative abundance of bacterial communities for the control (n=5) or exposed to selected bacteria (n=5/condition) oyster samples at the Phylum level
